# A Demographic History of a Prairie Vole (*Microtus Ochrogaster*) Breeding Colony (2004-2020)

**DOI:** 10.64898/2026.02.20.707040

**Authors:** Adele M H Seelke, Christina L Hung, Sabrina L Mederos, Sophia Rogers, Tiffany Lam, Lauren A Meckler, Karen L Bales

**Author notes:** Corresponding author: Adele M H Seelke PhD.

## Abstract

Prairie voles (*Microtus ochrogaster*) are highly social rodents that have become a valuable animal model for studying social attachment, pair bonding, parental care, and the neurobiological mechanisms underlying social behavior. In recent years, due in part to the publication of the prairie vole genome and deeper mechanistic understanding of their social behavior, prairie voles have become a more popular research model, especially for translational research. However, generating reliable and reproducible findings requires effective colony management, including thoughtful breeding strategies, consistent husbandry practices, and clear documentation. In this paper, we describe the demographic history of and husbandry techniques employed in our prairie vole breeding colony at UC Davis from 2004 to 2020. Well-organized and transparent colony management allows for the preservation of informative behavioral traits in prairie voles and strengthens the impact of the prairie vole model across behavioral and biomedical science.

## Introduction

Prairie voles (*Microtus ochrogaster*) are microtine rodents native to the central grassland regions of the United States ^1^. In the wild, they inhabit burrow systems that offer shelter and nesting areas for their offspring, often forming long-term family units ^2,3^. These social structures are characterized by behaviors uncommon in most rodents, such as social monogamy, biparental care, and selective aggression toward unfamiliar conspecifics after bonding with a mate ^4,5^.

In a research context, prairie voles are frequently studied to investigate the genetic, hormonal, and neural mechanisms underlying social attachment and pair bonding, where a pair-bond is a psychological construct that is present if individuals form enduring bonds with a single mate, show distress upon separation, and display preference for their partner over novel conspecifics ^6–10^. These behaviors are reliably replicated in laboratory conditions, where captive prairie voles also readily breed, adapt well to handling, and exhibit naturalistic social behaviors ^6,8^. Given this behavioral consistency across environments, prairie voles have become a premier model species for investigating the neurobiological and genetic mechanisms underlying social attachment and monogamy ^6,8,9^.

Over the last few decades, prairie voles have been instrumental in mapping out the roles of oxytocin and vasopressin systems in affiliative behaviors and attachment ^8,11–14^. The publication of the annotated prairie vole genome, along with the development of sophisticated genomic tools, has further enabled researchers to explore complex questions about gene regulation, neuroplasticity, and behavioral expression ^15–18^. Their relatively short lifespan of 2 – 12 months in the wild ^2^, combined with sexual maturation by postnatal days 45 – 60 ^19^ and frequent breeding cycles, allow for longitudinal and multigenerational studies within a feasible timeframe ^6,8^.

As the use of prairie voles in laboratory research continues to expand, it becomes increasingly important to maintain stable, well-characterized, and reproducible colony conditions ^6,8^. Well-documented colony husbandry is essential not only for ensuring continuity in long-term studies, but also for interpreting experimental variability across different research institutions ^20,21^. While wild populations offer key insights into natural behavior and ecology, laboratory colonies present an opportunity to study behavioral and physiological traits under standardized conditions, improving our ability to isolate specific genetic, environmental, and social variables. However, relatively little data are available on long-term reproductive patterns in captive prairie voles, especially when compared to more common laboratory rodents such as mice and rats.

The demographic structure of a captive prairie vole colony is shaped by numerous factors, including breeding strategies, housing arrangements, generational turnover, and overall colony management. Variables such as birth rates, survival rates, litter sizes, sex ratios, and weaning timelines are crucial metrics for understanding colony dynamics. These metrics can influence experimental design, interpretation of results, and overall animal welfare. Differences in housing protocols, environmental enrichment, and pairing strategies between laboratories may also contribute to subtle yet significant differences in behavioral outcomes. Early handling and parental care manipulations produce long-term and even intergenerational changes in social behavior and neuroanatomy ^22–26^. Enrichment and prior pairing history also affect social preferences, aggression, and other behaviors reinforcing the need to document these variables for cross-lab reproducibility ^8,27,28^ Documenting these factors is critical for standardizing practices, facilitating reproducibility, and optimizing animal husbandry for this model species.

The goal of the present study is to provide a comprehensive description of how prairie voles live and breed within our laboratory colony and describe specific trends in their reproduction. By detailing our methods for housing, pairing, and tracking colony demographics over time, we aim to create a reference framework for other researchers establishing or maintaining prairie vole colonies. This work also contributes to a broader understanding of how demographic variables may influence experimental outcomes, allowing for more accurate cross-lab comparisons and improved scientific rigor.

Our laboratory has maintained a colony of prairie voles continuously since 2004, utilizing standardized husbandry procedures throughout. Here, we present demographic information from 134 pairs of prairie voles that produced 2475 litters between the colony’s inception in 2004 and March 2020, when research was disrupted due to the Covid-19 pandemic. These data have been collected through the transcription of archival breeding records.

## Materials and Methods

### Housing and Husbandry

The prairie vole colony was established using voles from the colony of C. Sue Carter at the University of Illinois, Chicago, and was housed in the Psychology department vivarium on the campus of the University of California, Davis. The lights were maintained on a 14:10 light/dark cycle, with lights on at 6 AM and off at 8 PM. This long photoperiod encourages breeding ^29,30^. Room temperature was maintained between 68-74°F and humidity was monitored and generally found to be within 35-50%.

Breeding pairs of animals were housed in large cages (10½” W x 19” L x 6⅛” H) that contained 4.5-5 cups of Sani-Chips (PJ Murphy; Montville, NJ, USA) as bedding along with two cotton nestlets (Ancare; Bellmore, NY, USA). Non-breeding animals were pair housed with same-sex age-matched conspecifics in small cages (7½” W x 11½” L x 5” H) containing 2-2.5 cups of Sani-Chips bedding. All cages were topped with wire lids with spaces for food and water bottles (Ancare; Bellmore, NY, USA).

All animals were provided ad libidum oval pellet food (equivalent to LabDiet catalog #5326-3; custom-milled LabDiet oval pellet High Fiber Rabbit Feed). Additional small pellet High Fiber Rabbit Feed (LabDiet catalog # 5326; St. Louis, MO, USA) was provided daily for breeder pairs, daily for 10 days post-weaning for weanlings, and weekly for all animals during cage changes. Water was dispensed ad libidum to each cage with a heavy-duty 16 oz glass bottle topped with double-sealed stoppers and ball waterer drinking tubes (Ancare; Bellmore, NY, USA).

All cages and water bottles were changed once weekly. Lids were replaced and racks were sanitized bi-weekly.

### Breeder pair formation

Breeder pairs were formed with a focus on maintaining genetic diversity within the colony. Animals designated as breeders could not have grandparents in common and preferably did not have great-grandparents in common (maximum 3% - 4% relatedness coefficient) to avoid inbreeding. After animals reached sexual maturity at postnatal day 45 ^31^, a male and female were placed together in a breeder cage with two cotton nestlets, bedding, and extra small pellet food. The animals were monitored for aggression following pairing. In the rare cases where aggression occurred, the individuals were separated for at least 10 days and were either removed from contention or re-paired with different individuals. Once the breeder pairs began to produce litters, they were considered successfully paired and remained permanently housed together. If the rare case that a breeder pair failed to produce a litter in three months, the pair was either separated for future use or culled. If an individual breeder died, its mate was culled.

When warranted by experimental conditions, we employed a timed mating protocol adapted from Kenkel, 2019 ^32^. Sexually mature males and females were placed together in a large cage with an olfactory stimulus, small amount of soiled bedding from the male’s cage. The pairs are also given cotton nestlets for bedding material. Twenty-three hours later, voles were separated from each other using a clear plexiglass cage divider with small holes to prevent copulation. This still allowed visual, auditory, and olfactory stimulation. Two days later, the cage divider was removed, allowing for mating to occur. This day was considered as the day of conception, or embryonic day 0 (E0).

In efforts to increase genetic diversity, we also periodically imported prairie voles from outside institutions when our breeder colony arrived at a “genetic bottleneck” which we defined as being unable to generate new breeding pairs that did not share grandparents. These animal imports were negotiated between the Bales lab and other labs that maintain prairie vole colonies and usually involved importing 8-10 individuals that were then used to generate new breeding pairs. During the period of time covered by this study, we received imported prairie voles from colonies maintained at North Carolina State University in 2013 and Florida State University in 2016.

### Colony management

Researchers performed daily animal welfare checks. Daily checks consisted of recording high and low temperatures and humidity from the previous 24 hours, counting the number of animals in each cage and ensuring it matched the number on the cage identification card, checking food and water and refilling as necessary, and checking cages for flooding or excessively soiled bedding. Every animal in the colony was observed to ensure general good health and counted for a daily colony-wide census.

Cages containing breeding pairs were checked daily for newborn pups. When new litters were found, the date of birth was recorded. The number of pups was recorded at postnatal day 2 in order to ensure all pups had been delivered. Female prairie voles have six nipples, so are only able to nurse six pups at a time; if there were more than six pups present, the litter was culled down to six. Pups were weaned at postnatal day 20 ±1.

During weaning, the juvenile voles were sexed. Same-sex pairs were placed in small cages and assigned animal identification numbers (IDs). For 10 days post-weaning, weanlings were provided with extra small pellet food in addition to the standard large food pellets. Sexes were confirmed 7 – 10 days after weaning, when external genitalia became more easily distinguishable. We avoided individually housing animals due to the social nature of prairie voles. In cases with an odd number of weanlings, they were preferably housed in groups of three and only individually housed if there were no other options. If an animal needed to be housed individually, it was given a cotton nestlet for enrichment and every attempt was made to rehouse it with an age- and size-matched conspecific within a week. Unrelated animals housed together were distinguished using ear punches, which were noted on their cage cards. Following weaning, females were moved to a female-only room in the facility. Males were housed in either a male-only room or in a room that contained breeder pairs and virgin males housed in same-sex pairs.

The following naming convention was used for IDs: breeder pairs were numbered sequentially, and each litter was identified as offspring of a specific breeder pair. Individual animals within a litter were signified by a letter affixed to the end of the ID, with letters assigned in weaning order, beginning with males. For example, an animal with the ID 325-12A indicated the following information: the individual was born to pair 325, was a member of their 12^th^ litter, and was the first animal weaned from that litter. This naming scheme enabled easy identification of related animals, especially those offspring that share parents.

### Animal handling

Handling prairie voles presents unique challenges. Their tails are short and have a high risk of degloving, so picking them up by their tails is not advised. Likewise, prairie voles show more aggressive behavior towards humans than traditional laboratory rodents. Therefore, for both personal and animal safety researchers from our lab always wear a protective leather or garden glove to handle animals.

When handling is necessary, researchers use either a cupping or scruffing technique. The cupping technique involves directing the vole into a soft silicone cup (10.5 cm tall x 8 cm diameter) and covering the cup’s mouth with a gloved hand or carefully pinching part of the opening closed, which minimizes direct handling of the animal. In the scruffing technique, the researcher uses their non-dominant hand, protected by a glove, to pick up the vole by the skin above the shoulders. Scruffing allows for secure and precise animal handling, and it is used when researchers need to closely evaluate the body (e.g., sexing weanlings or monitoring health conditions).

Special care is used when handling animals in breeding cages, which can be either the adult mating pair or their young pups. Nursing prairie vole mothers typically have pups latched to their nipples by milk teeth ^33^, so when a nursing mother is transferred, the latched pups must be transferred with her. In this case, the researcher scruffs the mother and holds their free hand under the mother and litter, supporting the weight of the attached pups. When the maternal transfer is complete, any remaining unlatched pups are then transferred by being gently scooped into a gloved hand.

Nursing mothers are always transferred by scruffing rather than cupping, as early life handling and husbandry has profound long-term impacts on parents, pups, and pups’ subsequent offspring. Parents handled using the cup technique display more pup-directed behaviors compared to scruffed parents ^34^, while for pups, being handled with the cup technique results in less alloparental care in males, impaired pair bonding in females, and more anxiety-like behaviors in both sexes ^35^. Early life handling manipulations also have intergenerational consequences, with the offspring of cup-transferred pups showing less alloparental care than offspring of scruff-transferred pups ^26^. Together, this indicates that early-life husbandry techniques have both immediate and intergenerational, sex-specific consequences on behaviors relevant to pair bonding and biparental care.

### Archival breeder data

Data were compiled from archival breeding records, including cage cards, weaning logs, and mortality logs. The subjects presented here were chosen for analysis based on three factors: the completeness of their records, their presence on the main breeding colony protocol, and the fact that they did not undergo experimental manipulations at any point.

For each subject, we recorded their date of birth, date of death, sex, date of pairing with a mate, and the number of litters born to a breeder pair. For each litter, we recorded their date of birth, the number of pups born, and, where available, the sex ratio of the litter. From these data we calculated fertility over time, inter-litter interval, reproductive span, and effects of seasonality.

### Statistical Analyses

General linear regression analyses on annual and seasonal variation in litter size were performed using R version 4.4.0 using package *stats* version 4.4.0 ^36^. All other analyses, including unpaired t-tests, correlation, and linear regression were performed using Graphpad Prism version 10.5.0. In all cases, alpha was set at 0.05.

## Results

Data were compiled from 134 pairs of prairie voles that lived within the UC Davis Psychology Vivarium between 2004 and 2020. These 134 breeding pairs produced 2475 litters and a total of 11023 pups.

We started by examining the demographic characteristics of the breeding voles within our colony. Breeding pairs were formed, with few exceptions, at least 15 days after both males and females reached sexual maturity (after postnatal day 60). Female voles were paired at 91.7 ± 3.99 days (range 33-304 days), while males were significantly older, at 136.1± 7.13 days of age (range 31-363 days; t266 = 5.423, p < 0.0001; Fig 1A). Previous experience indicated that male voles remained fertile longer than females, so we chose to pair older males with younger females when establishing breeder pairs.

**Figure 1.**
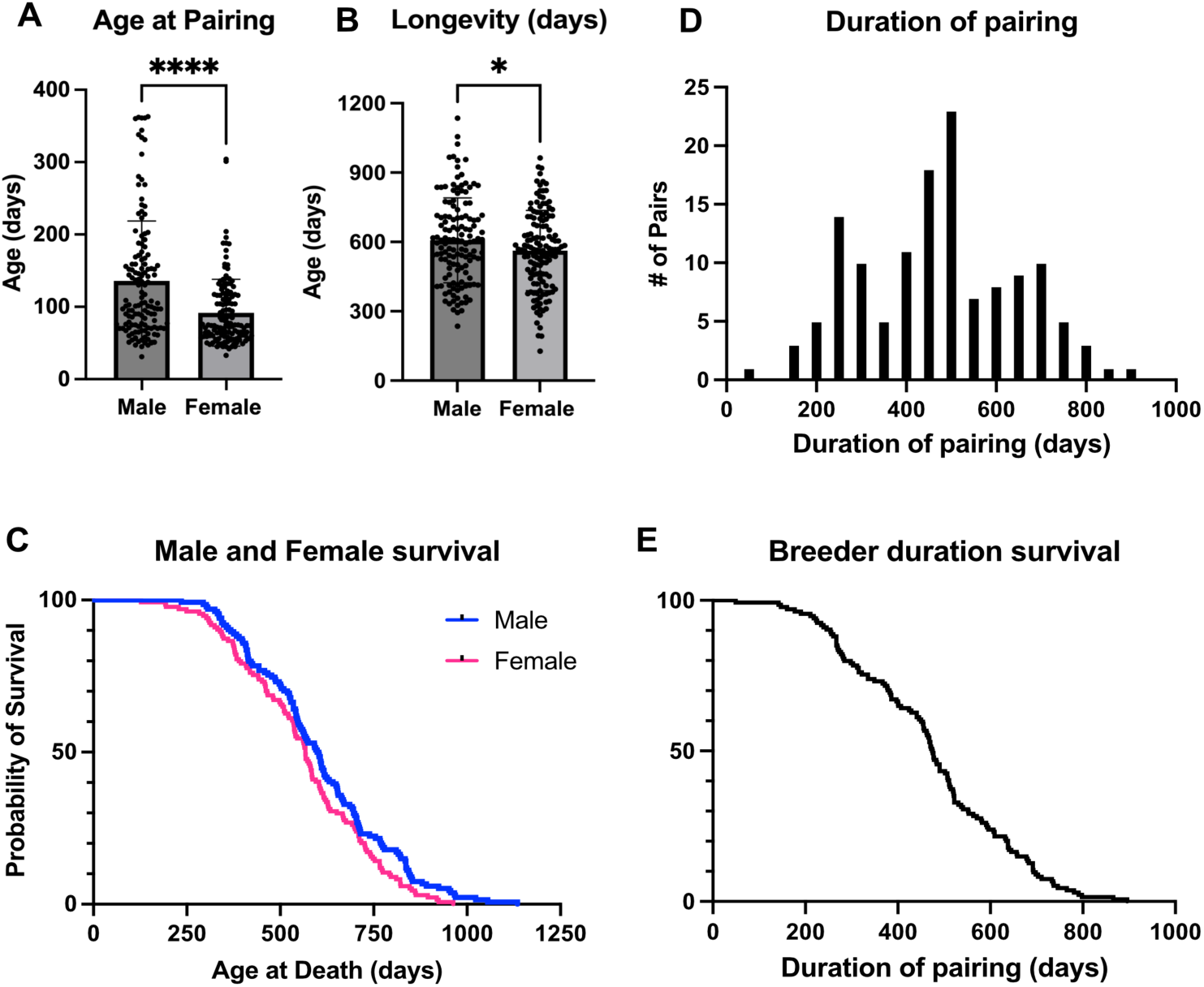
Demographic information about male and female prairie voles used as breeders. A) The mean age at pairing in days of male (left) and female (right) voles. The average age at pairing was significantly lower for females than males. B) The mean longevity in days of male (left) and female (right) prairie voles used as breeders. The average life span of males was significantly longer than females. C) A survival plot showing the longevity of male and female breeder voles in days. The x-axis shows an individual’s age at death, and the y-axis shows the proportion of subjects that were still alive at that age. A Mantel-Cox analysis revealed a significant difference between the survival curves of males and females, with the female curve shifted to the left relative to the male curve. D) A frequency distribution of the duration prairie vole breeding pairs. The x-axis shows the duration of pairing in days while the y-axis shows the number of pairs with a specific duration. The average pair duration was 471.1 ± 14.8 days. E) A survival plot showing the duration of pairing of male and female breeders. The x-axis shows the duration of the pair, and the y-axis shows the proportion of breeders that were still paired at that age. **** - p < 0.0001, * - p < 0.05

The mean life span of breeding male voles was 607.2 ± 15.82 days, while the mean lifespan of breeding females was significantly shorter at 562.8 ± 14.96 days (t266 = 2.035, p = 0.0428; Table 1; Fig 1B). A Mantel-Cox survival analysis revealed a significant difference between the survival curves of males and females (C^2^(1, N = 134) = 4.523, p = 0.0334; Fig 1C). The average duration of pairing, defined as the time from pair formation until the death of one pair member, was 471.1 ± 14.8 days, ranging from 49 – 896 days (Fig 1D,E). We should emphasize that the lifespan of voles described by these data is not necessarily representative of a full lifespan; instead, it reflects the reproductive lifespan of prairie voles.

**Table 1.**

|  | **Age at Pairing (days)** |  | **Age at Cull (days)** |  | **Duration of Pairing (days)** |
| --- | --- | --- | --- | --- | --- |
|  | *Male* | *Female* | *Male* | *Female* |  |
| **Mean** | 136.1 | 91.7 | 607.2 | 562.8 | 471.1 |
| **s.d.** | 82.5 | 46.2 | 183.2 | 173.2 | 171.4 |
| **Min** | 31 | 33 | 235 | 127 | 49 |
| **Max** | 363 | 304 | 1135 | 963 | 896 |

We next examined demographic characteristics of the pups born to the established breeding pairs within our colony. We calculated the inter-litter interval (ILI), defined as the number of days from the birth of one litter until the birth of the next litter. This was calculated for both every individual litter (N = 2475 litters; Fig 2A) and for each breeder pair (N = 134 breeder pairs; Fig 2B). The mean ILI across the entire population was 24.07 ± 0.04 days with a range of 19 – 140 days (Fig 2A; Table 2), while the mean ILI for individual breeder pairs was 25.6 ± 0.3 days with a range of 21.8 – 42.6 days (Fig 2B). We then calculated the mean number of pups born to each litter, which was 4.47 ± 0.03 pups with a range of 1 – 10 pups in a single litter (Fig 2C). We next calculated that the mean number of litters born to breeder pairs was 18.5 ± 0.6 with a range of 2 – 38 litters (Fig 2D).

**Figure 2.**
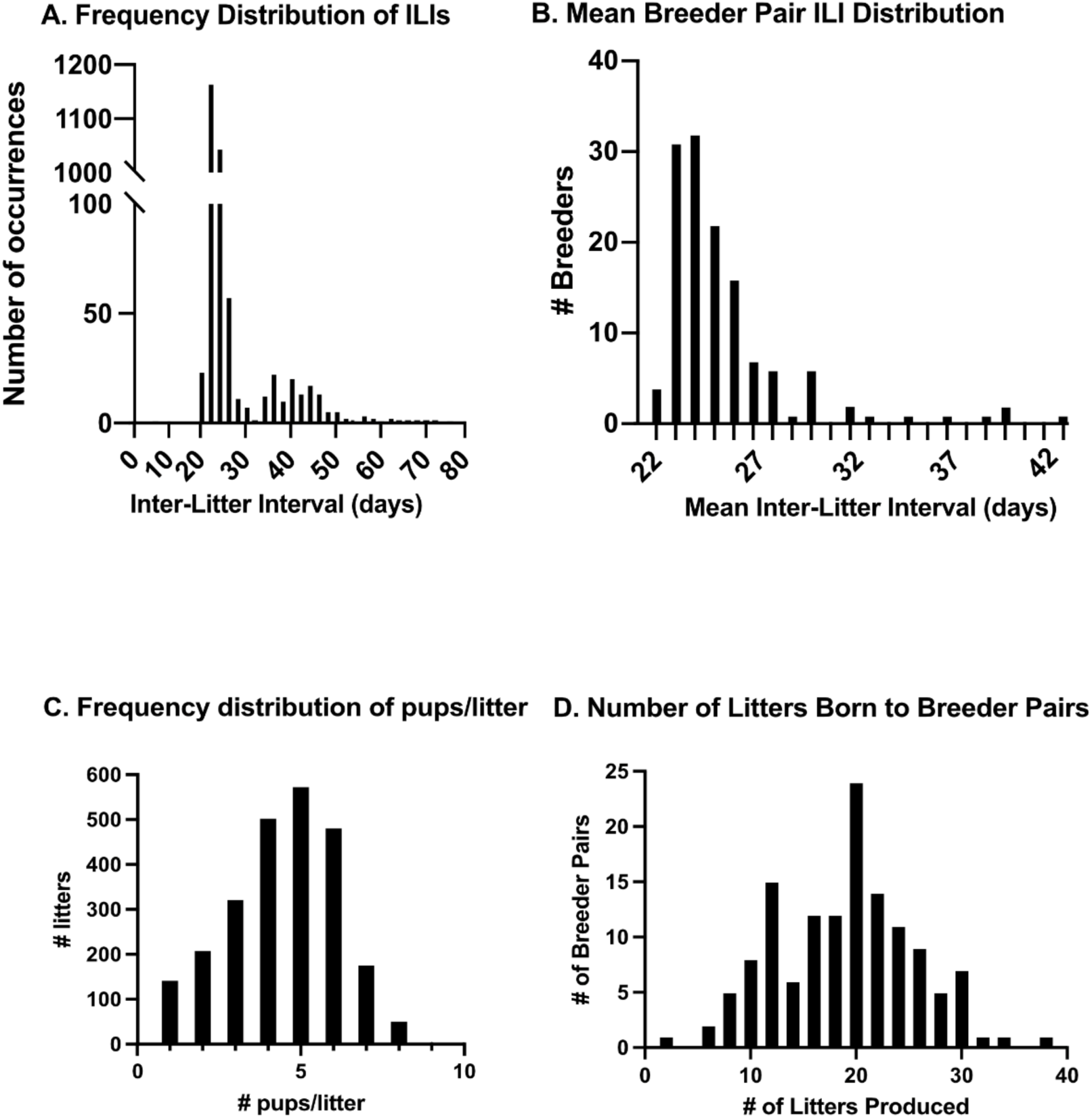
Demographic information about the litters generated by prairie voles breeder pairs. A) Histogram demonstrating the inter-litter interval (ILI) frequency distribution for all 2475 litters included in this study. There was an extremely pronounced peak in the histogram in the bins containing ILIs of 22 – 25 days, and the mean ILI was 24.07 ± 0.04 days. B) We calculated the average ILI of each breeder pair to examine variation between breeder pairs. The histogram shows the frequency distribution of mean ILIs across breeders, with a peak in the bins containing ILIS of 23 – 25 days. We also observed several breeder pairs with extremely long mean ILIs, demonstrating the variability that existed between individual breeder pairs. C) Histogram showing the frequency distribution of the number of pups born in a litter (x-axis) and the number of litters that contained a given number of pups (y-axis). There is a peak in the frequency distribution at 5 pups per litter. D) Histogram showing the frequency distribution of the number of litters produced by a single breeder pair. The x-axis shows the number of litters, and the y-axis shows the number of breeder pairs that produced a given number of litters. There is a peak in the frequency distribution at 20 litters.

**Table 2.**

|  |  |
| --- | --- |
| **Total # breeder pairs** | 134 |
| **Total # litters** | 2475 |
| **Mean inter-litter birth interval (days)** | 24.07 ± 0.04 |
| **Mean # litters/ breeder pair** | 18.47 |
| **s.d. # litters** | 6.55 |
| **Max # litters** | 38 |
| **Min # litters** | 2 |
| **Total # pups** | 11023 |
| **Mean pups/litter** | 4.46 |
| **s.d. pups/litter** | 1.69 |
| **Max pups/litter** | 10 |
| **Min pups/litter** | 1 |

We then began to explore the sources of variability within litters born to our established breeding pairs, including the relationship between litter size and sex ratio of the pups. We analyzed a subset of the archival data that contained information about the sex ratio of pups within a litter (1292 out of 2474 litters), which included 5602 pups (2779 male, 2823 female) out of a total 11023 pups within the entire dataset. The mean number of male pups per litter was 2.151 ± 0.035, while the mean number of female pups per litter was 2.185 ± 0.037 (t2582 = 0.6746, p = 0.50, ns).

We next examined how the sex ratio of pups changes with litter size. We grouped litters by the number of pups, then for each litter with X number of pups we determined the proportion of litters that contained Y number of males. For example, for litters that contained 4 pups (X = 4; Fig 3D), we calculated that 6.5% of these litters contained 0 males, 21.3% contained 1 male, 40.9% contained 2 males, 25.1% contained 3 males, and 6.2% contained 4 males. Figure 3 (A-H) shows how the number of male pups in a litter changes as the size of the litter increases. Each graph indicates the number of pups in a litter, the total number of litters of that size, and the number of male and female pups represented in the data. The distribution of sex ratios appears to approach normality with the increase in number of subjects, centered around an approximately 1:1 male to female sex ratio.

**Figure 3.**
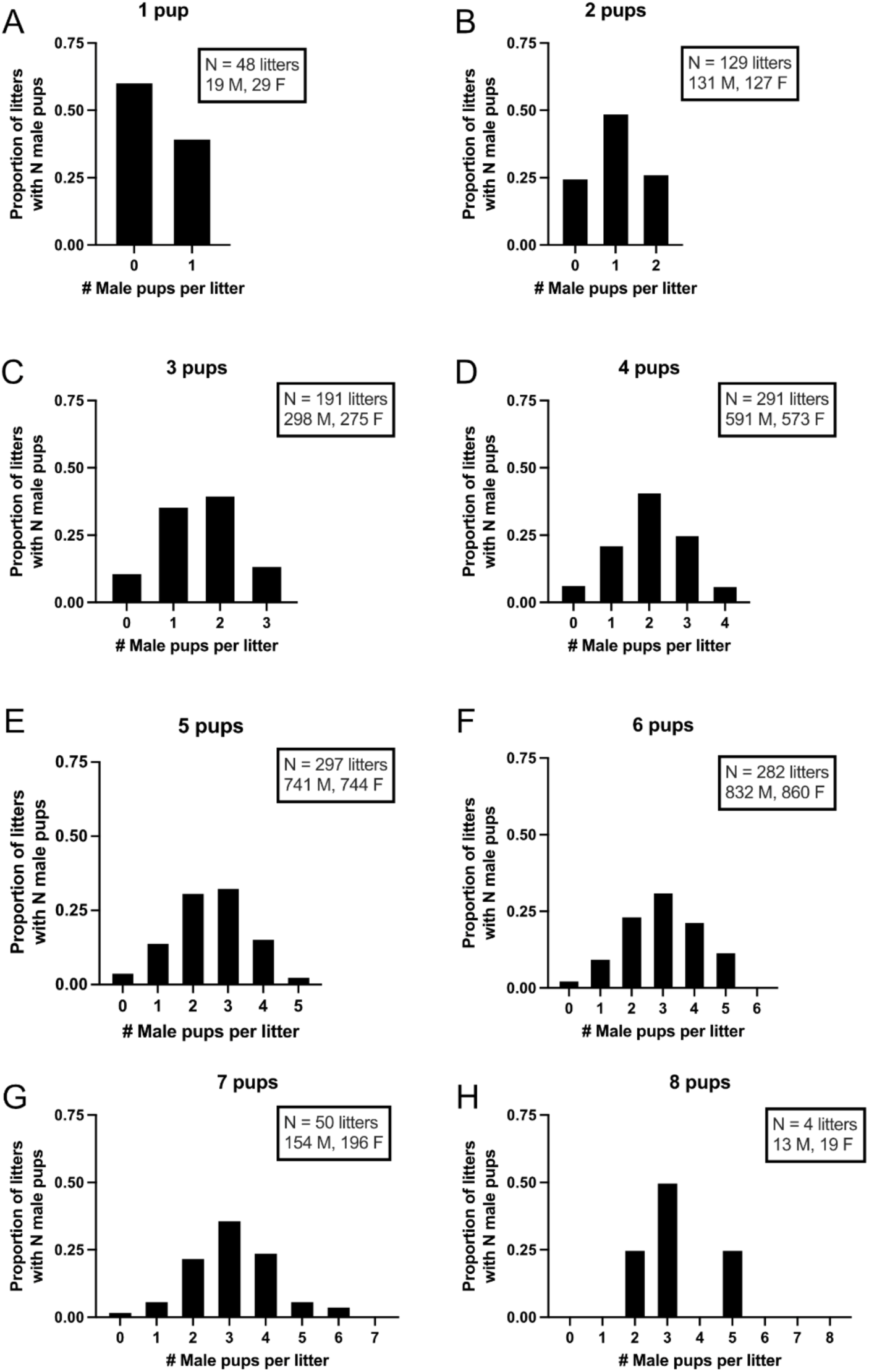
The ratio of male to female pups in litters of different sizes. The x-axis depicts the number of male pups in a litter, and the y-axis depicts the proportion of litters that contain that number of male pups. In each graph, the number of litters included in the analysis, as well as the total number of male and female pups in all the litters, is shown. A) The proportion of male pups in litters that contained 1 pup. B) The proportion of male pups in litters that contained 2 pups. C) The proportion of male pups in litters that contained 3 pups. D) The proportion of male pups in litters that contained 4 pups. E) The proportion of male pups in litters that contained 5 pups. F) The proportion of male pups in litters that contained 6 pups. G) The proportion of male pups in litters that contained 7 pups. H) The proportion of male pups in litters that contained 8 pups. As the both the number of pups per litter and the number of litters included in the analysis increases, the distribution of pup sex ratios appears to approach normality.

We also looked for annual and seasonal variation in the number of pups per litter born from 2005 – 2019, which excluded years when the colony had not been operational for the entire year (Fig 4). Examining birth month across years, there was also no significant effect of seasonality on litter size (general linear regression, F11, 2448 = 0.9174, p = 0.5224; Fig 4A). There was no significant effect of birth year on litter size (general linear regression, F1, 2458 = 0.6109, p = 0.4345; Fig 4B).

**Figure 4.**
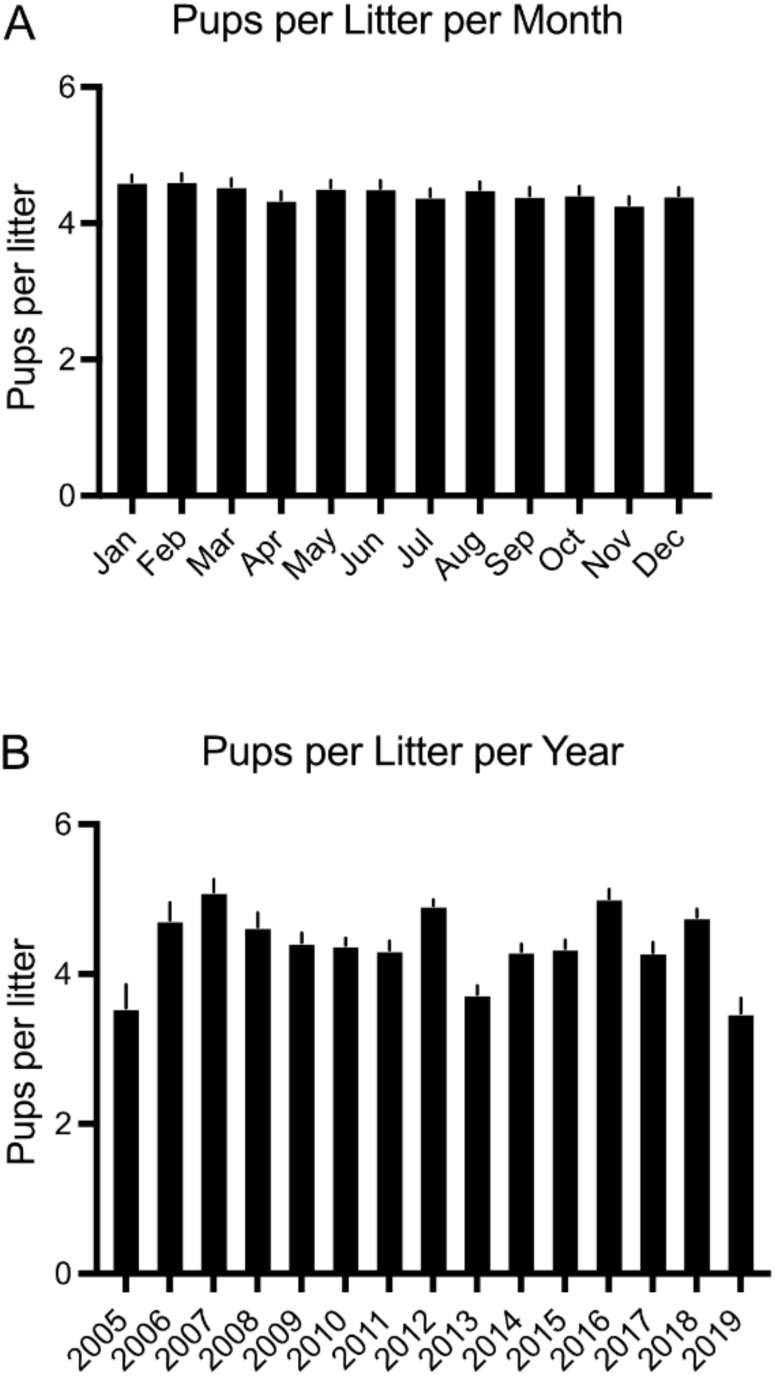
Seasonal and annual variation in the number of pups born per litter. Years without litter information available for the complete calendar year (2004, 2020) were excluded. A) Bar plots showing the number of pups born per litter in each month during the years 2005 – 2019. Error bars represent 1 standard error measurement. There is no significant difference between months, indicating an absence of seasonal variation in vole fertility in this colony. B) Bar plots showing the number of pups per litter born each year from 2005 – 2019. Error bars represent 1 standard error measurement. There are no significant differences between years, indicating an absence of annual variation in vole fertility in this colony.

As previous analyses did not find any significant annual or seasonal variation in litter size, we decided search for other sources of variation within our breeding colony, beginning with an examination of how breeding changes across the lifespan. We first examined how inter-litter interval (ILI) changed with breeding experience. As the number of litters born to a breeder pair increases, so do ILI duration (r = 0.1659, p < 0.0001; Fig 5A) and proportion of ILIs with durations above the mean (r = 0.7668, p < 0.0001; Fig 5B). The exception to this is the latency from initial breeder pair formation until the birth of the first litter, where 32.1% of first litters are born later than would be expected by the average ILI. Another analysis demonstrated a significant negative correlation between ILI duration and number of pups born (r = -0.1986, p < 0.0001; Fig 5C), indicating that as ILI duration increases, the number of pups born decreases.

**Figure 5.**
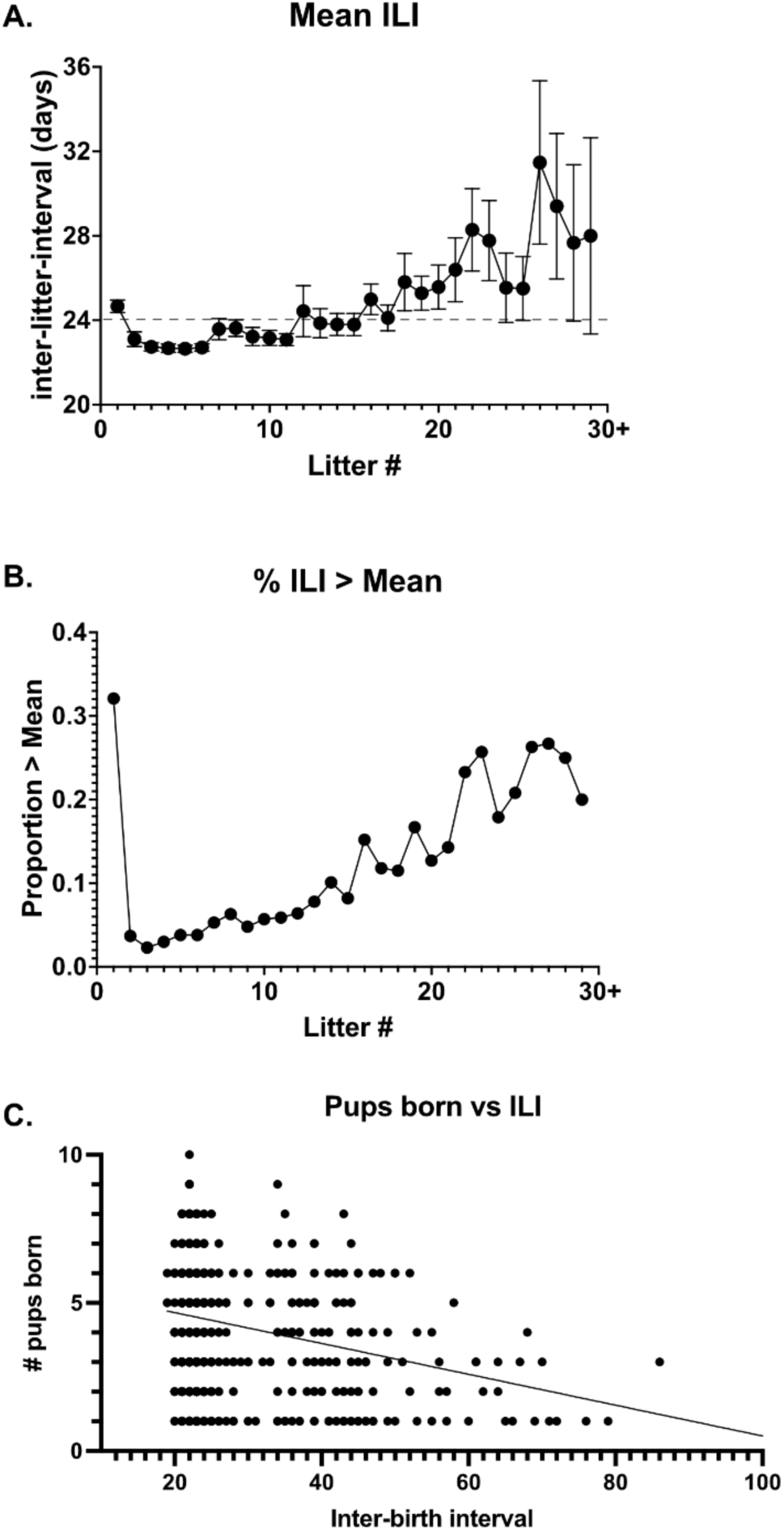
Changes in inter-litter interval (ILI) across time. The ILI was defined as the number of days between the birth of one litter and the birth of the subsequent litter. A) For each breeding pair, we calculated the ILI between each litter and then calculated the mean ILI for each litter number. At the number of litters born increased, the ILI increased as well. B) The mean ILI was calculated for all litters, and the proportion of litters born with an ILI greater than the mean ILI was determined. The ILI of the first litter was defined as the time between pairing and the birth of the first litter, and this mean duration was longer than that of the following litters. C) There is a significant negative correlation between ILI (x-axis) and number of pups born (y-axis), indicating that as the time between the birth of litters increases, the number of pups born decreases.

We next examined how fertility changed across the lifespan of breeding voles, beginning with the relationship between parental age and characteristics of a breeder pair’s first litter. The father’s age at the birth of the first litter ranged from 59 – 386 days, with a mean of 160.5 ± 7.2 days. The mother’s age at the birth of the first litter ranged from 69 – 258 days, with a mean of 117.1 ± 4.0 days. There was no effect of father’s age on number of pups or time to birth of first litter, and no effect of mother’s age on time to birth of first litter. There was a small but significant effect of mother’s age on the number of pups born in the first litter (F1,131 = 6.188, p = 0.0141). Across the lifespan, as the number of litters born to a breeder pair increased, the number of pups born per litter significantly decreased (F1,2467 = 248.4, p < 0.0001; Fig 6A). However, there was a strongly significant positive correlation between the duration of pairing and number of litters produced (r = 0.9331, p < 0.0001; Fig 6B). The survival plot in Fig 6C shows the proportion of breeding pairs (y-axis) that produced a given number of litters (x-axis).

**Figure 6.**
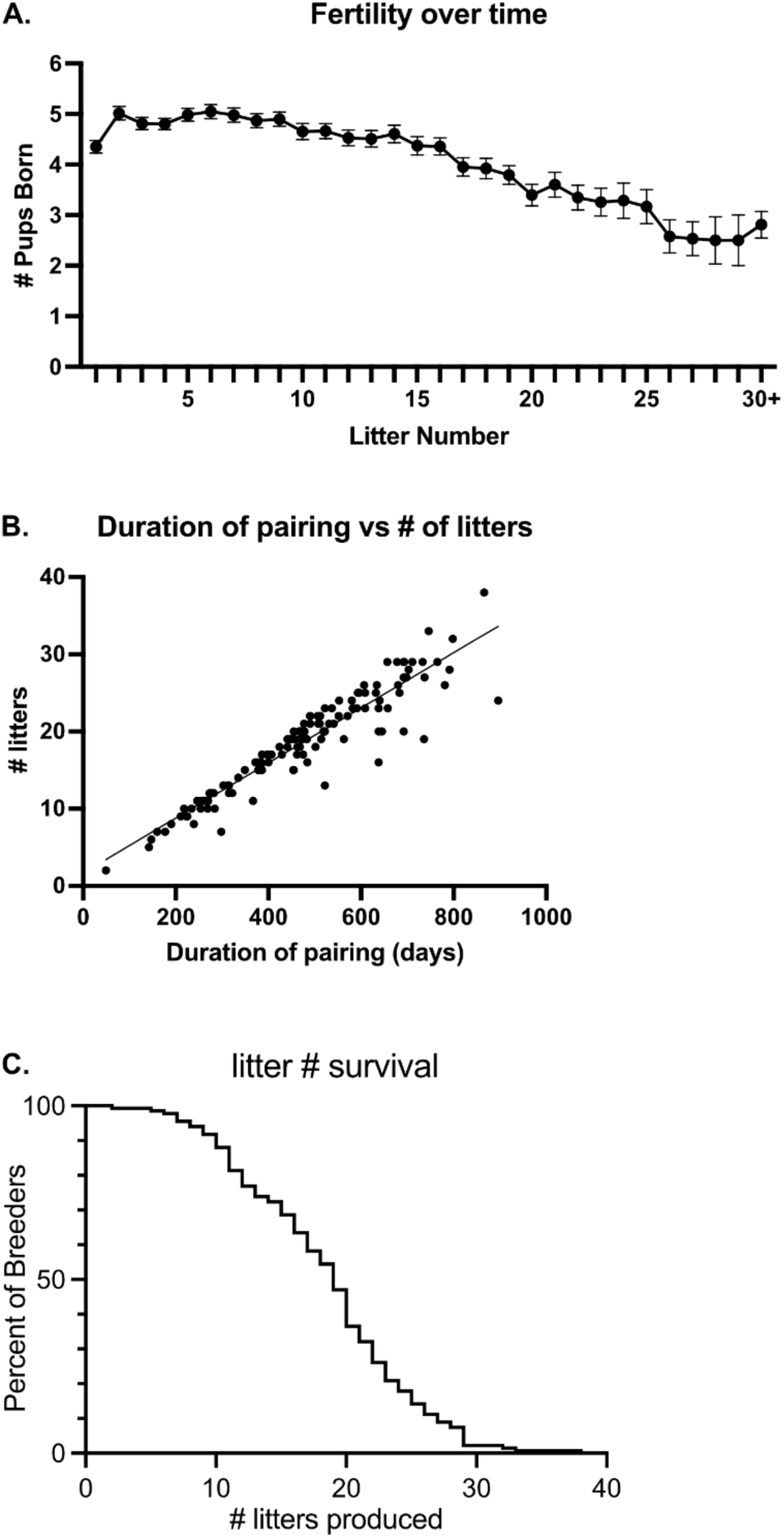
Changes in breeder fertility over time. A) The average number of pups born (y-axis) to a given litter (x-axis). As the number of litters born to the breeder pair increases, the number of pups born in a litter significantly decreases, demonstrating a decrease in fertility over time. B) There is a significant positive correlation between the duration of pairing of a specific breeder pair (x-axis) and the number of litters produced by that breeder pair (y-axis). The fact that the correlation is linear indicates that voles remain fertile and capable of producing litters across their entire lifespan. C) A survival plot showing the number of litters produced by a breeding pair (x-axis). The y-axis shows the proportion of breeding pairs that produced a given number of litters.

## Discussion

Our current husbandry framework was established in 2004 under the direction of Dr. Karen Bales. These management practices reflect a continuation of practices originally created and refined by Dr. C. Sue Carter, and the collective experience of several other long-standing colonies at other universities. The implementation of consistent husbandry conditions and standardized breeding protocols has been critical to the long-term stability and productivity of our colony. We hope that by highlighting our methodology and colony demographics here, we can demonstrate a possible lab setup for current and future prairie vole researchers.

Breeding prairie voles can present a set of unique challenges compared to other more commonly used laboratory rodents. Their formation of long-term pair bonds can preclude immediate re-pairing ^28^ and often limits the number of potential genetic combinations within a colony. Some breeding techniques common in lab mice and rats (polygamous mating, male rotation, etc.) cannot be directly applied to prairie voles without risks to animal welfare. If a breeder pair in our colony was unsuccessful, i.e. no pups were produced or no litters were weaned after approximately three months or 3 – 4 breeding cycles, we separated them for an extended period (around 2 – 3 weeks) before attempting to re-pair the individuals with new mates to preserve every lineage for as long as possible.

Nonetheless, such re-pairing is uncommon, as once a successful pair bond is established, the pair will typically remain together for the duration of their lives. Our prairie vole breeding pairs are usually stable once formed, lasting 471.1 days on average in our colony (Table 1). Each pair produced an average of 18.5 litters throughout their lifetime with 4 – 5 pups per litter (Table 2). Within our colony, we found no significant seasonal variation in litter size (Fig 4A), unlike the increased spring litter sizes found in wild prairie voles ^37^. Although litter size decreases significantly over time, this decrease is only by 1 – 2 pups on average (Fig 6A). This consistency across bloodlines differs from laboratory mice, where litters may have 2 – 12 pups but size varies widely by genetic strain (The Jackson Laboratory).

Prairie vole females experience postpartum estrus ^38^, which occurs within 24 – 48 hours of parturition ^39^. Prairie vole litters typically gestate for 20 – 23 days ^40^, slightly longer than the typical 19 – 20 day gestation of a lab mouse pregnancy ^41^. As the mean birth interval between litters observed in our colony is 24.1 days (Table 2), this suggests that postpartum estrus is usually productive in our breeders. By contrast, lab mice experience postpartum estrus around 14 – 28 hours after parturition and have fewer successful fertilizations in postpartum estrus compared to during the regular estrus cycle ^41^. This greater consistency in our vole colony allows for some level of predictability in litter birth dates and subsequent animal availability for experimental recruitment.

In order to maintain the health and naturalistic behaviors of wild-type prairie voles, it is essential that the colony remains outbred. Maintaining sufficient genetic diversity requires careful management and periodic introduction of new animals, a challenge our laboratory has addressed by coordinating with other established prairie vole colonies to receive new animals as needed (typically every two to three years). This ensures our colony obtains an ongoing influx of novel genetic material.

The genetic diversity in our outbred colony is reflected in its phenotypic diversity. Although there is generally little variation in prairie vole physical appearance, some of our colony’s bloodlines appear to have subtle differences from each other, such as more grey-toned vs yellow-toned underbelly fur, darker brown vs tawnier body fur, and so forth. Behaviorally, we have long recognized the natural variation that can be seen in pup-directed parental behaviors ^22^, thus we characterize the parental care of every breeder pair in our colony as part of our standard husbandry practices. Variation in parental care received can have lasting impacts on the offspring’s behavior and neurobiology (Seelke et al, 2016; Tabbaa et al, 2017; Rogers & Bales, 2019; Perkeybile et al, 2019; Bottom et al, 2020; Rogers et al, 2021), making records of parenting “style” for each breeder pair a valuable resource. The natural variability of prairie voles provides validity to translational behavioral and physiological research using this species.

## Conclusion

Prairie voles are a unique translational model that is key to studying a variety of topics, including social attachment, pair bonding, and affiliative behavior, as well as how these processes are altered by early life adversity, genetic variation, and drug exposure. Prairie voles are an especially valuable species for these questions compared with traditional laboratory rodents because they are outbred, socially monogamous, form pair bonds, and display biparental care. The work conducted in prairie voles has demonstrated that the formation of pair bonds can alter neural circuits and neuropeptide pathways, primarily oxytocin, vasopressin and dopamine, in ways that provide the foundation for lasting attachment and parental investment ^12,42^.

Prairie voles are increasingly common models for studies examining social communication and attachment deficits, including developmental and neuroendocrine studies testing pharmacological interventions that target social behavior ^8,9^. More recently, with the sequencing of the prairie vole genome, genetic and molecular tools have been developed that allow researchers to perform detailed manipulations on the level of neural circuits ^15,43^. Further research in this area could be vital to understanding the natural variation in oxytocin, vasopressin and dopamine systems and expand our ability to test causal relationships between genes, neural circuits and complexity of social behavior.

Ensuring that prairie vole colonies are well managed is essential for preserving the unique social behavior of this valuable translational model. Careful attention to breeding practices, housing conditions and lineage tracking enhances transparency and allows other researchers to accurately reproduce findings. Strong colony management ultimately ensures that vole-based research remains reliable, interpretable, and scientifically impactful.

## Acknowledgements

We gratefully acknowledge the work performed by former graduate students and staff associated with the Bales lab, including Cindy Clayton DVM, Mrs. Jessica Bond, Anita Stone PhD, Caroline Hostetler PhD, Allison Perkeybile PhD, Trenton Simmons PhD, and Forrest Rogers PhD. We also thank the Campus Veterinary Services of the University of California, Davis, especially Dr. Rhonda Oates DVM. And we extend a special thanks to Dr. C. Sue Carter, who is one of the mothers of prairie vole research. We gratefully extend the trail that she helped blaze.

## Funding Statement

During the time period covered by this manuscript, the prairie vole colony was supported by funding from the following organizations:

The National Institutes of Health

5 R01 MH108319 – Molecular and Neural Networks Underlying Social Attachment
4 R01 HD071998 – Effects of Chronic Intranasal Oxytocin
5 R21 HD060117 – Effects of Early Experience on Somatosensory Systems in Voles

The National Science Foundation

ADVANCE Fellows Award (0437523): Effects of Early Experience on Parental Behavior in Voles

University of California, Davis Office of Research

